# Host and environment shape the giant clam-associated photosymbiont community

**DOI:** 10.64898/2026.08.07.743467

**Authors:** John Bennedick Quijano, Kuselah M. Tayaban, Jake Ivan P. Baquiran, Gabriella Juliane Maala, Jeremiah Noelle C. Requilme, Sherry Lyn G. Sayco, Roger G. Dolorosa, Patrick C. Cabaitan, Cecilia Conaco

## Abstract

Giant clams are some of the largest bivalve molluscs. They form a vital partnership with Symbiodiniaceae dinoflagellates that supply most of their energetic requirements. However, the factors that shape giant clam-associated photosymbiont communities remain unknown. Here, we profiled Symbiodiniaceae communities using ITS2 metabarcoding in eight giant clam species (*Hippopus hippopus*, *H. porcellanus*, *Tridacna crocea*, *T. derasa*, *T. gigas*, *T. maxima*, *T. noae* and *T. squamosa*) from 11 sites across the Philippine archipelago. Symbiodiniaceae community structure was shaped by an interplay between giant clam host and environment. Most giant clams were dominated by members of a single symbiont genus, with *Cladocopium* as the most prevalent, followed by *Durusdinium* and *Symbiodinium*. However, giant clam hosts also exhibited flexibility in their symbiotic partners that was evident across sites. Differences in giant clam-associated symbiont communities may contribute to differences in holobiont function and adaptability to variable environments. These findings deepen our understanding of giant clam-Symbiodiniaceae associations, offering a framework for predicting how giant clams may be affected by increasingly stressful reef conditions and, more importantly, informing strategies to improve mariculture and conservation practices.

## INTRODUCTION

Giant clams (Cardiidae: Tridacninae) are the largest living bivalves found mostly in Indo-West Pacific coral reefs (Rosewater 1965; bin Othman et al. 2010; Hui et al. 2016; Neo et al. 2017). These iconic clams provide food and habitat for other reef organisms and contribute to primary production through their partnership with photosynthetic dinoflagellates from the family Symbiodiniaceae (Klumpp and Griffiths 1994; Cabaitan et al. 2008; Neo et al. 2015). Among the 12 extant giant clam species, eight, including *Tridacna gigas*, *T. squamosa*, *T. noae*, *T. maxima*, *T. crocea*, *T. derasa*, *Hippopus hippopus* and *H. porcellanus*, are found within the Coral Triangle, including most parts of the Philippines (bin Othman et al., 2010; Neo et al., 2017; Tan et al., 2022). Populations of these giants, however, are dwindling due to excessive overfishing owing to their appeal as ornamentals and food source. Overfishing has resulted in local extinction of *T. gigas* and *T. derasa* in the Philippines (Juinio et al. 1989; bin Othman et al. 2010; Neo and Todd 2013; Juinio-Meñez et al. 2025). The International Union for Conservation of Nature (IUCN) Red List of Threatened Species lists *T. gigas* as critically endangered and *T. derasa* as endangered (Neo and Li 2024a, 2024b). As a response to plummeting stocks, various conservation efforts have focused on enhancing giant clam aquaculture and restocking on depleted reefs (Gomez and Mingoa-Licuanan 2006; Waters et al. 2013; Mies et al. 2017a).

Like corals and other photosymbiotic invertebrates, giant clams host Symbiodiniaceae (LaJeunesse et al. 2018). These microalgal symbionts are acquired exclusively from the environment (Mies et al. 2017b) and hosted extracellularly in specialized tubules within the giant clam (Norton et al. 1992; Poo et al. 2020). Symbiodiniaceae provide up to about 97% of the clam’s energy requirements, delivering necessary metabolites such as glucose, amino acids, glycerol, and sterols (Norton et al. 1992; Ishikura et al. 1999; Mies al. 2017a; Hambleton et al., 2019; Uchida et al., 2026). In return, the host provides nutrients that support photosynthesis (Armstrong et al. 2018; Pang et al. 2022) and offers an environment that can enhance symbiont photosynthetic performance and reduce UV stress (Holt et al. 2014; Rossbach et al. 2020). Diverse Symbiodiniaceae genera, including *Symbiodinium* (formerly Clade A), *Cladocopium* (formerly Clade C), and *Durusdinium* (formerly Clade D), associate with giant clams (Baillie et al. 2000; Pappas 2017; Lim et al. 2019; Guibert et al. 2020). This photosymbiont diversity varies across species and growth stages and is influenced by environmental conditions and geographic locations (DeBoer et al. 2012; Ikeda et al. 2017). Symbiodiniaceae also play an important role in shaping characteristics of the host, including growth rate and development (Long et al. 2021), and resistance to heat stress-induced bleaching (Pappas 2021; Mies et al. 2025).

Despite the importance of Symbiodiniaceae to giant clam survival, little is known about their diversity and the factors governing their association with giant clams. In this study, we explored the diversity and composition of Symbiodiniaceae in eight giant clam species across selected sites in the Philippines and examined the factors driving symbiont community structure. Our findings revealed that Symbiodiniaceae communities are shaped primarily by giant clam species but also influenced by local conditions. This pattern suggests that the giant clam-Symbiodiniaceae partnership is driven mainly by the host but is flexible enough to allow adjustments based on the environment. The study improves our understanding of the giant clam-Symbiodiniaceae symbiosis, from the general diversity of photosymbionts associated with giant clams to the specific Symbiodiniaceae types linked to particular hosts and environments. Furthermore, these findings can be used to inform giant clam mariculture practices, as early introduction of a suitable population of Symbiodiniaceae may be a crucial step in enhancing growth and survival of giant clams and could potentially improve the outcomes of giant clam restocking.

## METHODS

### Collection sites

Eight species of giant clams (*Tridacna gigas*, *T. derasa*, *T. maxima*, *T. squamosa*, *T. noae*, *T. crocea*, *Hippopus hippopus* and *H. porcellanus*) were sampled between 2018-2020 from 11 sites representing five marine biogeographic regions of the Philippines (Figure 1a, STable 1). Calaguas is in the North Philippine Sea (NPS), Samar is in the South Philippine Sea (SPS), Bohol and Camiguin are in the Visayan Sea (VS), Semirara Marine Hatchery Laboratory (SMHL) and Palawan are in the Sulu Sea (SS), and Bolinao, Hundred Island National Park (HINP), Batangas, and Apo Reef Natural Park (ARNP) are in the West Philippine Sea (WPS) (Aliño and Gomez 1994). Giant clams from Calaguas and ARNP represent natural wild stocks, as these sites do not have history of restocking prior to this work. On the other hand, all giant clams from Bolinao and SMHL were hatchery-reared and grown out in ocean nurseries. Most *T. gigas* individuals that were sampled across sites are restocked hatchery-cultured giant clams originating from the Silaqui ocean nursery at Bolinao, Pangasinan (Gomez and Mingoa-Licuanan 2006; Mingoa-Licuanan and Gomez 2007), with a few juvenile individuals (shell length ca. 35 cm) from Camiguin derived from natural recruitment (Requilme et al. 2021). Other giant clam species from the other sites are wild stocks but may include a few hatchery-reared giant clams, as there are records of multi-species restocking at these sites (Gomez and Mingoa-Licuanan 2006; Mingoa-Licuanan and Gomez 2007), although no long-term monitoring information is available.

**Figure 1.**
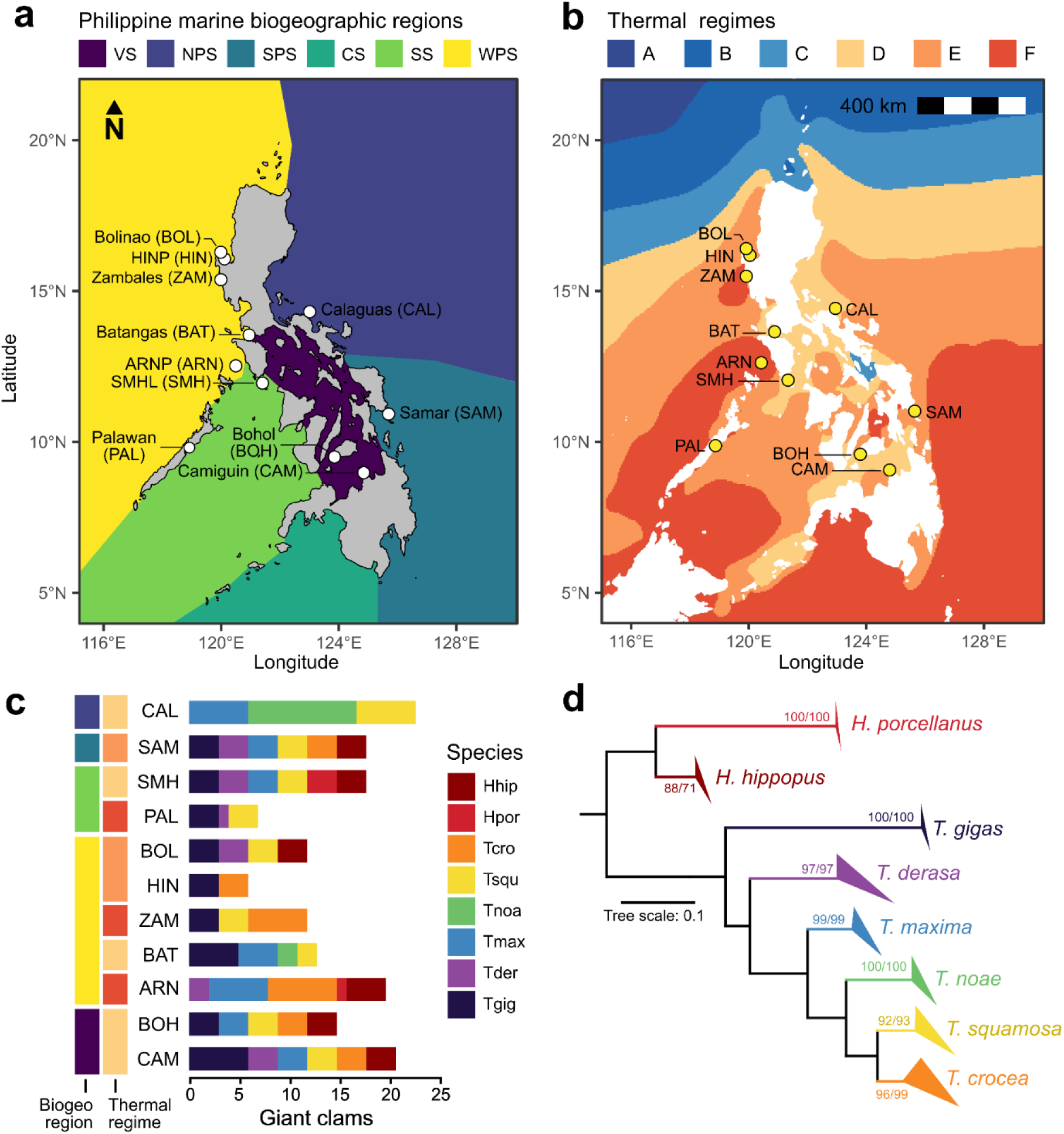
Sampling of giant clams across Philippine biogeographic regions and thermal regimes. (**a**) Sampling sites (CAL, Calaguas; SAM, Samar; SMH, SMHL; PAL, Palawan; BOL, Bolinao; HIN, HINP; ZAM, Zambales; BAT, Batangas; ARN, ARNP; BOH, Bohol; CAM, Camiguin) within marine biogeographic regions (VS, Visayan Sea; NPS, North Philippine Sea; SPS, South Philippine Sea; CS, Celebes Sea; SS, Sulu Sea; WPS, West Philippine Sea). (**b**) Thermal regimes identified using k-means clustering of SST data. (**c**) Giant clam species sampled per site, biogeographic region, and thermal regime. (**d**) Maximum-likelihood dendrogram based on the COI marker confirming species identity of samples. Branch support values are shown as SH-aLRT/UFBoot. Scale bar represents branch length.

### Sample collection

Sample collection was performed by SCUBA divers. Each giant clam individual was photographed *in situ* and an initial species identification was made based on visual observation. A small non-lethal clipping (approximately 0.5-1 cm^2^) was taken from the mantle of each giant clam using surgical scissors and placed into a 5 ml tube with ambient seawater. Tissues were transferred into salt-saturated DMSO-EDTA (SSDE) buffer for storage and were kept at 4°C until DNA extraction. Collections were conducted with permission from the Department of Agriculture Bureau of Fisheries and Aquatic Resources (Gratuitous Permit Nos. 0169-19, 0188-20, 0189-20, 0190-20) and the Palawan Council for Sustainable Development (Wildlife Gratuitous Permit 2018-32).

### DNA extraction

A small piece of mantle tissue was taken from each sample using sterile scissors and homogenized with a plastic pestle. Total DNA was extracted using the Qiagen DNeasy Plant Mini Kit (Qiagen, Germany) according to the manufacturer’s instructions. DNA quality was examined by agarose gel electrophoresis and quantified using a microvolume spectrophotometer (Nanodrop 2000, Thermofisher, USA).

### COI gene sequencing, phylogeny, and haplotype analyses

Giant clam host identity was verified by amplifying a fragment of the COI gene using specific primers (STable 2) (Folmer et al. 1994; Nuryanto et al. 2007; Lizano and Santos 2014). PCR products were visualized by agarose gel electrophoresis and submitted to Macrogen, South Korea, for Sanger sequencing. Forward and reverse COI sequences were assembled and checked manually for miscalls using Molecular Evolutionary Genetics Analysis 11 (MEGA 11). Quality-filtered sequences were aligned with giant clam sequences from GenBank using MAFFT L-INS-i 7.475 (Katoh and Standley 2013). Phylogenetic inference was done using maximum likelihood implemented in IQ-TREE (Nguyen et al. 2015; Trifinopoulos et al. 2016) with the model HKY+F+R2 model selected based on BIC values computed using ModelFinder (Kalyaanamoorthy et al. 2017). Clade support was determined using Shimodaira-Hasegawa approximate likelihood ratio test (SH-aLRT, Guindon et al. 2010) and ultrafast bootstrap approximation (UFBoot, Hoang et al. 2018) based on 1,000 bootstrap replicates.

Haplotype relationships were inferred using statistical parsimony networks in pegas (Paradis 2010). For each species, COI sequences were aligned, trimmed to remove ambiguous base positions, and collapsed into unique haplotypes per species. Haplotype networks were constructed using the haploNet function, which infers relationships among haplotypes based on the number of mutational steps separating sequences. Haplotype diversity (Hd) and nucleotide diversity (π) were calculated using functions implemented in the same package.

### ITS2 metabarcode sequencing and analysis

The ITS2 region was amplified using the primer pair ITSintfor2 (LaJeunesse and Trench 2000) and ITS2-reverse (Coleman et al. 1994). Libraries were generated using the Herculase II Fusion DNA Polymerase Nextera XT Index Kit V2 (Agilent Technologies, Santa Clara, CA, USA). Metabarcode sequencing was done at Macrogen, South Korea, on the Illumina MiSeq platform with version 3 chemistry to generate 300 bp paired-end reads. Resulting sequences were analyzed using the SymPortal framework through a local run (Hume et al. 2019). Raw paired fastq.gz files were submitted to SymPortal and subjected to quality control (QC) using mothur 1.39.5 (Schloss et al. 2009), BLAST+ suite of executables (Camacho et al. 2009) and minimum entropy decomposition (MED) (Eren et al. 2015). Symbiodiniaceae-related ITS2 sequences were identified and the most common and abundant sequences (>= 200 reads) were classified as defining intragenomic variants (DIVs). DIVs were then grouped into ITS2 type profiles, which serve as proxies for putative Symbiodiniaceae genotypes or strains. Assignment of ITS2 type profiles minimizes over-estimation of community diversity due to intragenomic diversity, i.e., the presence of multiple ITS2 sequences in a single Symbiodiniaceae genotype (Hume et al. 2019).

### Symbiodiniaceae community analysis

Shannon diversity of ITS2 type profiles among groups were compared using Kruskal-Wallis Test (Kruskal and Wallis 1952) and pairwise comparisons were made using Dunn’s test (Dunn 1964). *P*-values from pairwise comparisons were adjusted using Benjamini-Hochberg false discovery rate correction (Benjamini and Hochberg 1995). Bray-Curtis dissimilarity (Bray and Curtis 1957) was computed from square-root transformed counts of ITS2 sequences (Oksanen et al. 2022), then non-metric multidimensional scaling (NMDS) plots (Kenkel and Orlóci 1986) were generated to visualize community structure across groups. Homogeneity of dispersion among clusters was assessed using PERMDISP2 (betadisper) (Anderson 2006). Differences in Symbiodiniaceae community composition among groups were analyzed using permutational multivariate analysis of variance (PERMANOVA; adonis2) (Anderson 2001) and pairwiseAdonis (Martinez Arbizu 2020), with each group tested in separate models using marginal tests (9,999 permutations).

The relationship between host haplotype diversity and ITS2 type profile diversity was assessed using a Spearman rank correlation test. Host genetic distances per species based on COI sequences were calculated using the Kimura two-parameter model implemented in the R package ape (Paradis and Schliep 2018). Indicator DIVs were identified using Indicator Species Analysis in the indicspecies package (duleg = TRUE), using an indicator value (IndVal) ≥ 0.5 and *P*-value ≤ 0.05 based on 999 permutations (De Cáceres and Legendre 2009; De Cáceres et al. 2010).

### Phylosymbiosis and cophylogeny analysis

Phylosymbiosis was assessed by evaluating topological congruency and distance congruency (Li et al. 2025). Topological congruency was examined by comparing the giant clam COI phylogeny with photosymbiont dendrogram constructed using unweighted pair group method with arithmetic mean (UPGMA) based on Bray-Curtis dissimilarity (Sokal and Michener 1958). Tree congruence was quantified using the normalized Robinson-Foulds metric (Robinson and Foulds 1981; Lim and Bordenstein 2020), with significance assessed (9,999 permutations) using the RFmeasures function (Mazel et al. 2018). Distance congruency was tested by evaluating the correlation between host phylogenetic distance and symbiont Bray-Curtis dissimilarity. Spearman correlation was assessed using a Mantel test (9,999 permutations) (Mantel 1967).

Cophylogeny between giant clams and *Symbiodinium*, *Cladocopium*, and *Durusdinium* communities was assessed using the Procrustean approach to cophylogeny (PACo) (Balbuena et al. 2013). Giant clam phylogenetic distances were derived from COI sequences, while ITS2 type profiles were represented by UniFrac dissimilarity. PACo was performed using symmetric Procrustes superimposition, with global model fit quantified as m^2^, the sum of squared residuals, and expressed as R^2^ = 1 - m^2^ (Perez-Lamarque and Morlon 2024). Statistical significance of the cophylogenetic signal was assessed using 9,999 permutations, where *P* < 0.05 and R^2^ values closer to one reflecting strong cophylogenetic congruence (Perez-Lamarque and Morlon 2024). The contribution of each host-symbiont association to the overall fit was calculated using paco_links() function, which computes jackknifed squared residuals for each link, where lower residual values reflect stronger cophylogenetic congruence and higher values indicate weaker associations. Differences in jackknifed squared residuals among giant clam species were tested using a Kruskal-Wallis test. For significance testing, Dunn’s post-hoc pairwise comparisons were performed with Benjamini-Hochberg (BH) adjustment. Tanglegrams were made using the cophylo() function of the phytools package (Revell 2024).

### Thermal regime characterization

Marine thermal regimes across the Philippines were characterized by k-means clustering (Lloyd 1982) of daily sea surface temperature (SST) from 1985-2025, encompassing both seasonal and interannual temperature fluctuations to capture broad thermal patterns. Annual mean SSTs were computed for each thermal regime. Thermal stress exposure within each regime was quantified using DHW, which represents the accumulation of positive temperature anomalies over a 12-week period derived from CoralTemp SST data. Bleaching-level stress (4 ≤ DHW < 8 °C-weeks) and mortality-level stress (DHW ≥ 8 °C-weeks) were identified using the maximum DHW recorded per year. These metrics are proxies for sustained thermal stress, particularly during the El Niño-Southern Oscillation (ENSO) years when elevated solar irradiance and thermal anomalies are common in the region (Muñiz-Castillo et al. 2019; Torres et al. 2021). Daily SST and degree heating week (DHW) data were obtained from NOAA Coral Reef Watch at 5-km spatial resolution (Skirving et al. 2020) through the ERDDAP server (Wilson et al. 2020).

### Data processing and visualization

Datasets were cleaned, visualized and explored using RStudio (RStudio Team 2020) with the tidyverse suite of packages (Wickham et al. 2019), all within the R programming language (R Core Team 2023).

## RESULTS

### Giant clam diversity

A total of 165 giant clam mantle samples, including 32 *T. gigas*, 28 *T. maxima*, 29 *T. squamosa*, 25 *T. crocea*, 19 *H. hippopus*, 13 *T. noae*, 15 *T. derasa,* and four *H. porcellanus*, were collected from 11 sampling sites across five Philippine marine biogeographic regions (Figure 1a) and three thermal regimes (Figure 1b; SFigure 1a). Thermal regime D (27.8-29.0°C annual mean SST) encompasses Batangas, SMHL, Bohol, Calaguas, and Camiguin. Thermal regime E (28.1-29.4°C annual mean SST) includes Bolinao, HINP, and Samar. Thermal regime F (28.4-29.6°C annual mean SST) covers ARNP, Palawan, and Zambales (SFigure 1b). Patterns of Degree Heating Week (DHW), which reflect thermal stress exposure, broadly aligned with the thermal regimes (SFigure 1c).

The number of giant clam species that were sampled varied across sites. Six species were collected in Camiguin and Samar, five in ARNP and Bohol, four in SMHL, Bolinao and Batangas, three in Calaguas, Palawan and Zambales, and two in HINP (Figure 1c). The best represented species was *T. gigas*, which was collected from nine sites where restocking of hatchery-cultured individuals had previously been conducted (Gomez and Mingoa-Licuanan 2006; Mingoa-Licuanan and Gomez 2007). *Tridacna squamosa* was collected at eight sites, *T. maxima* and *T. crocea* at seven sites, *T. derasa* and *H. hippopus* at six sites, and *T. noae* and *H. porcellanus* at two sites. Identities of field-collected samples were verified using maximum likelihood (ML) analysis of COI gene sequences, which recovered 8 monophyletic groupings with high bootstrap support (≥85% SH-aLRT or ≥95% UFBoot) (Figure 1d and SFigure 2).

### Giant clam COI haplotypes

Genetic diversity within each species was estimated based on COI haplotypes (STable 3). The number of detected COI haplotypes (H) ranged from 1 to 20 per species (SFigure 3), with the greatest number of haplotypes seen in substrate-boring species, *T. crocea* and *T. maxima*. *Hippopus hippopus, T. crocea, and T. maxima* exhibited higher haplotype diversity (Hd ≥ 0.7), while *T. squamosa*, *T. noae*, and *T. gigas* and *H. porcellanus* had lower diversity (Hd ≤ 0.5) (STable 3). Most species had one or two dominant haplotypes and low nucleotide diversity (π) across sequences, with the exception of *T. crocea*, which did not have a single dominant haplotype but showed greater nucleotide diversity across sequences.

### Diversity of giant clam-associated Symbiodiniaceae

Sequencing of the ITS2 region yielded 18,173,186 contiguous amplicon sequences from 165 giant clam individuals, with a mean count of 110,140.52 (±21,200.87) amplicons per library (STable 4 and SFigure 4). After quality-checking and filtering, 15,396,367 (mean 93,311.32 ±18,807.07) sequences were analyzed on Symportal (STable 4 and SFigure 4). This analysis revealed 461 ITS2 sequences affiliated with four Symbiodiniaceae genera (STable 5) that were categorized into 33 ITS2 type profiles (STable 6).

The most abundant Symbiodiniaceae genus associated with Philippine giant clams was *Cladocopium* (61.77%), followed by *Durusdinium* (25.47%), and *Symbiodinium* (12.76%) (Figure 2a). *Gerakladium* was also present at negligible levels (<0.001%). *Cladocopium* was represented by 193 ITS2 sequences, with the most abundant being C93a, C1, C66, and C1c. *Durusdinium* also had 193 sequences dominated by D5, D4, D9b, and D1. *Symbiodinium* had 74 sequences with A3 and A6b as most abundant (Figure 2b, STable 5). C93a, D5, D4, and A3 were detected in around 60% of the giant clams (Figure 2c). Together, the ITS2 sequences defined 33 unique ITS2 type profiles, five for *Symbiodinium*, 17 for *Cladocopium*, and 11 for *Durusdinium* (SFigure 5).

**Figure 2.**
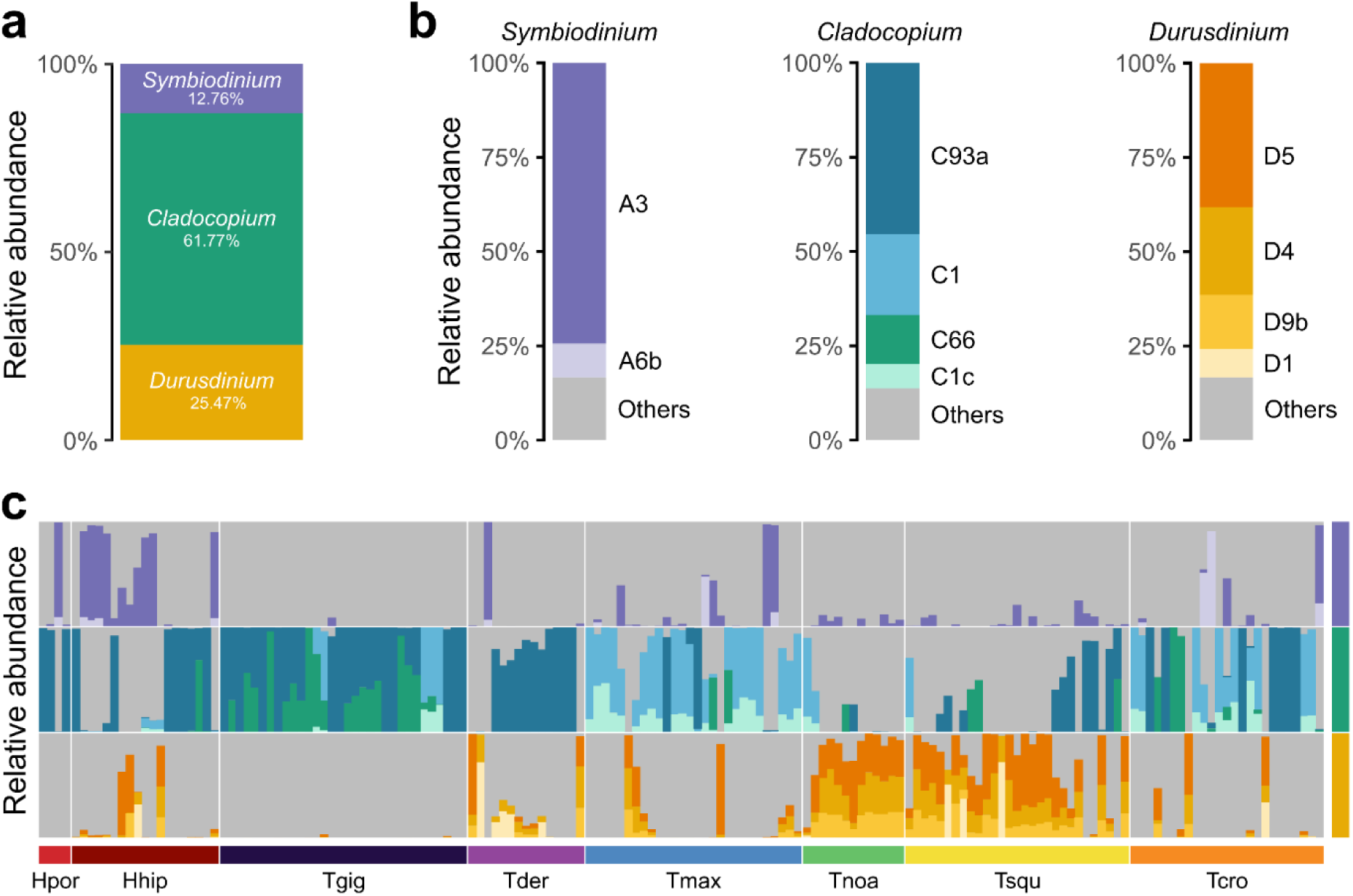
Symbiodiniaceae associated with Philippine giant clams. (**a**) Total relative abundance of Symbiodiniaceae genera. (**b**) Relative abundance of major ITS2 sequences within each Symbiodiniaceae genus (<1% total relative abundance grouped as “Others”). (**c**) Relative abundance of major ITS2 sequences (colors as in b) associated with individuals of each giant clam species (bottom color bar). Color bar on the right indicates symbiont genera (colors as in a).

Shannon diversity of Symbiodiniaceae based on ITS2 type profiles varied among giant clam species (Kruskal-Wallis, *H*(7) = 48.26, *P* < 0.001; SFigure 6a) but did not differ significantly across sampling sites, biogeographic regions, or thermal regimes (SFigure 6b-d). *Tridacna gigas* had the lowest symbiont diversity, with 84% of individuals harboring only a single ITS2 type profile. *Hippopus porcellanus* also showed relatively low ITS2 type profile diversity, although low sample size (n = 4) for this species limits inference (SFigure 7). Notably, COI haplotype diversity showed a weak positive correlation with diversity of Symbiodiniaceae ITS2 type profile (Spearman’s ρ = 0.21, *P* = 0.006).

### Drivers of photosymbiont community structuring

To assess the drivers of symbiont community structuring in giant clams, we examined Symbiodiniaceae ITS2 sequence composition across host species, sampling sites, biogeographic regions, and thermal regimes. We found that symbiont community structure differed significantly across these groups (Figures 3a-d; PERMANOVA; Table 1), with species explaining the greatest proportion of variation, followed by sampling sites, biogeographic regions, and thermal regimes. Multivariate dispersion differed significantly among species, biogeographic regions, and thermal regimes, but not among sampling sites (Figures 3a-d; PERMDISP; Table 1).

**Table 1.** PERMANOVA and PERMDISP results comparing Symbiodiniaceae composition among groups (9,999 permutations).

| Group | PERMANOVA |  |  | PERMDISP |  |
| --- | --- | --- | --- | --- | --- |
|  | R<sup>2</sup> | F | P | F | P |
| Species | 0.33243 | 11.169 | 0.001 | 4.0825 | < 0.001 |
| Sampling site | 0.25116 | 5.1651 | 0.001 | 1.4078 | 0.1815 |
| Biogeographic region | 0.12411 | 5.6681 | 0.001 | 7.4317 | < 0.001 |
| Thermal regime | 0.08925 | 7.9378 | 0.001 | 18.805 | < 0.001 |

At genus level, we observed that *T. gigas, T. derasa, T. maxima,* and *T. crocea* were predominantly associated with *Cladocopium*, while *T. noae* and *T. squamosa* tended to be dominated by *Durusdinium*. *Symbiodinium* was prevalent in *H. hippopus*, though some individuals of this species were dominated by *Cladocopium* (Figure 3e).

**Figure 3.**
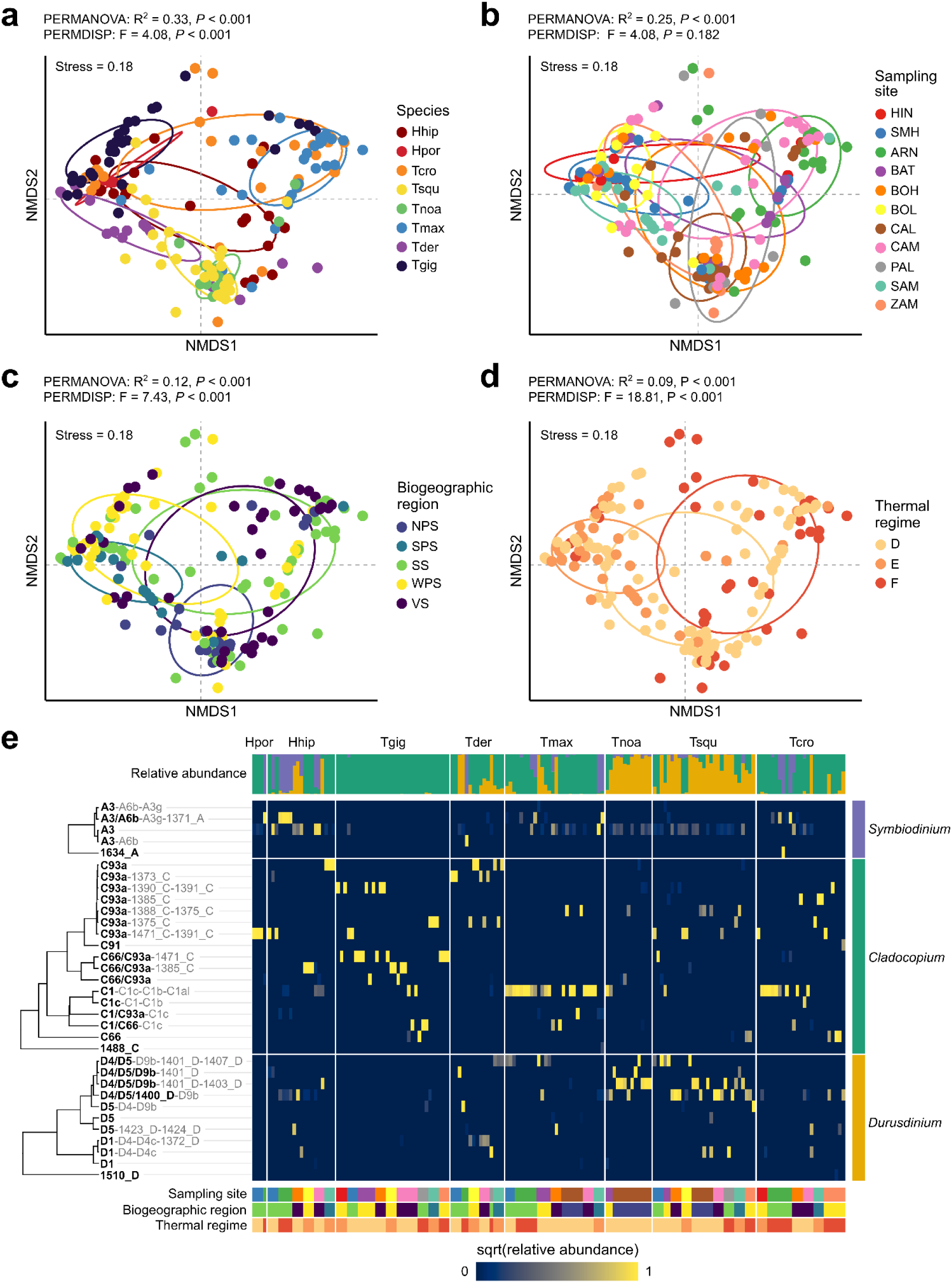
Symbiodiniaceae community composition among groups. NMDS ordination plots (stress = 0.18) showing Symbiodiniaceae ITS2 sequence composition among giant clam species (**a**), across sampling sites (**b**), biogeographic regions (**c**), and thermal regimes (**d**). Major statistics from PERMANOVA and PERMDISP are shown above each ordination. Points represent individual samples and are colored by group. Ellipses indicate group dispersion (confidence level = 0.5). (**e**) Relative abundance of ITS2 type profiles across all giant clam individuals (yellow, high; blue, low). Trees show average hierarchical clustering of ITS2 type profiles based on UniFrac distance. Major DIVs are highlighted in bold. The color bar on the right indicates symbiont genus. The top color bar indicates relative abundance of each ITS2 type profile (colored by genus) per individual. The bottom color bars indicate sampling sites, biogeographic region, and thermal regimes (colors as in a-d).

Detailed comparisons of ITS2 sequences among giant clam species revealed significant differences in photosymbiont communities, except between *H. hippopus* and *H. porcellanus*, and between *T. squamosa* and *T. noae* (SFigure 8a). This is supported by differences in shared ITS2 type profiles among giant clam species (Figure 3e, SFigure 9a). Indicator species analysis further identified defining intragenomic variants that characterized each giant clam species (SFigure 10a). *Symbiodinium* A3 was associated with *H. hippopus*. *Cladocopium* C66 was associated with *T. gigas,* C1c, C1, C1al, and C1b with *T. maxima*, and C93a, 1391_C, and 1471_C with *H. porcellanus*. *Durusdinium* D4, D5, D9b, 1401_D, and 1403_D were indicative of *T. noae*, D1 and 1372_D were associated with *T. derasa*, and 1400_D with *T. squamosa*.

Pairwise comparisons of ITS2 sequences among sampling sites showed compositional difference between most sites (SFigure 8b). The ARNP community, in particular, was distinct from all other sites. In addition, no ITS2 type profiles were shared across all sampling sites (SFigure 9b). Indicator species analysis showed that *Symbiodinium* A3, A6b, A3g, and 1371_A were significantly associated with ARNP, C*ladocopium* 1375_C with Samar, 1385_C with Bolinao, *Durusdinium* D1 and D4c with Palawan, 1407_D with Batangas, 1400_D with Bohol, and *Cladocopium* 1388_C and *Durusdinium* 1403_D and 1401_D with Calaguas (SFigure 10b).

Symbiodiniaceae communities differed significantly between biogeographic regions (SFigure 8c), except between Sulu Sea and Visayan Sea. Only three ITS2 type profiles were shared across all biogeographic regions, while a total of 13 were shared between the Sulu Sea and Visayan Sea (SFigure 9c). Indicator species analysis showed that *Symbiodinium* A6b, A3g, and 1371_A were significantly associated with the Sulu Sea, *Cladocopium* C93a and 1375_C with the South Philippine Sea, *Durusdinium* 1400_D with the Visayas Sea, and *Cladocopium* 1388_C and *Durusdinium* D9b, D5, D4, 1403_D and 1401_D with the North Philippine Sea (SFigure 10c).

Symbiodiniaceae communities were also distinct across thermal regimes (SFigure 8d). While ten ITS2 type profiles were shared among all regimes, an equal number were unique to individual regimes (Figure 3e, SFigure 9d). Indicator species analysis showed that *Durusdinium* D9b, D5, D4, and 1401_D were associated with regime D, *Cladocopium* C93a was associated with regime E, and *Symbiodinium* A6b, A3g, and 1371_A, and *Cladocopium* C66, were associated with regime F (SFigure 10d).

### Symbiodiniaceae community shifts in *Tridacna gigas*

To examine shifts in giant clam photosymbiont communities upon long-term translocation to different reef environments, we leveraged the availability of *T. gigas* individuals that had been restocked at various sites around the Philippines since the 1980s. These giant clams were sourced from hatchery-cultured individuals reared at the Silaqui ocean nursery in Bolinao, Pangasinan. Majority of the sampled *T. gigas* individuals shared haplotype G1. Some individuals from Batangas, Palawan, Samar, and Bohol belong to distinct but closely related haplotypes (Figure 4a).

**Figure 4.**
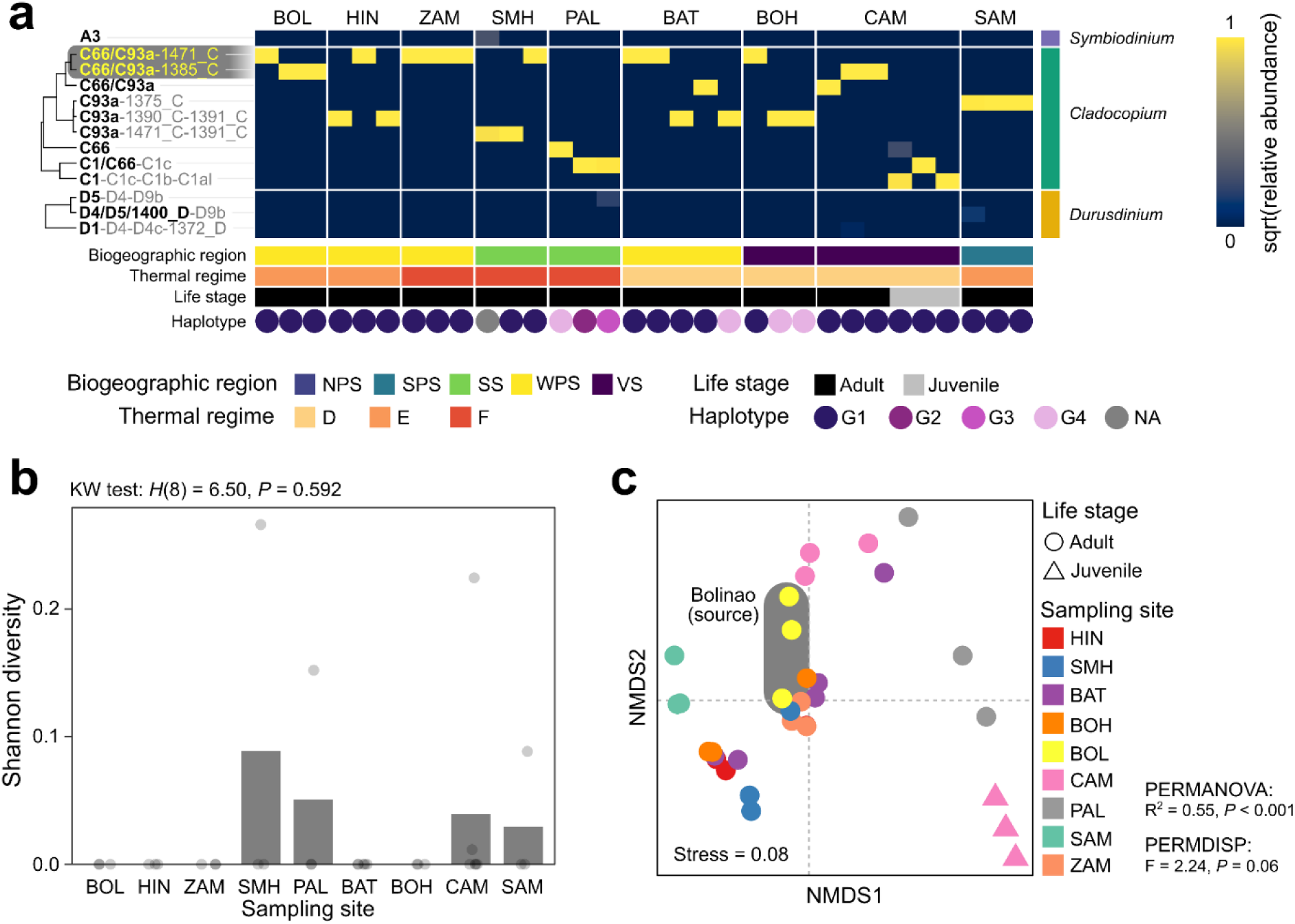
Symbiodiniaceae associated with translocated *T. gigas*. (**a**) Relative abundance of ITS2 type profiles across *T. gigas* individuals (yellow, high; blue, low) grouped by Symbiodiniaceae genus. Trees show average hierarchical clustering of ITS2 type profiles based on UniFrac distance. ITS2 type profiles highlighted in gray indicate those present in Bolinao (source). Major DIVs are highlighted in bold. The bottom color bars indicate sampling sites (colors as in b), biogeographic regions, thermal regimes, and life stage. Colored circles denote haplotype. (**b**) Mean ITS2 type profile diversity across regions. Kruskal-Wallis test results are displayed above. (**c**) NMDS ordination plot (stress = 0.08) showing Symbiodiniaceae composition in *T. gigas* across sampling regions (indicated by colors) and life stage (indicated by shapes). Individuals from the source site are highlighted in gray. Major statistics from PERMANOVA and PERMDISP are shown beside the plot.

The diversity of *T. gigas*-associated ITS2 type profiles did not differ significantly across sites (Figure 4b). Although global community comparison indicated significant site-driven structuring of Symbiodiniaceae composition (PERMANOVA: R^2^ = 0.55, *P* < 0.001), pairwise comparisons between the source and translocation sites were not significant after false discovery rate correction (Figure 4c; STable 7). Community dispersion also did not differ significantly across sampling sites (PERMDISP: F = 2.24, *P* = 0.06).

*Tridacna gigas* from the source site were dominated by ITS2 type profiles with C66/C93a as the major DIV. Most restocked *T. gigas* individuals at other sites hosted the same or closely related symbionts, still with C66/C93a or C93a as major DIVs (Figure 4a). Notably, *T. gigas* from Palawan and the juveniles from Camiguin hosted C66, C1/C66, or C1 symbionts, which belonged to a separate cluster from the other *T. gigas*-associated *Cladocopium* (Figure 4a).

#### Phylosymbiosis and cophylogeny in giant clams

Correlation between host COI genetic distance and Symbiodiniaceae community dissimilarity revealed a weak but significant signal of phylosymbiosis (Mantel test: Spearman r = 0.167, *P* < 0.001; 9,999 permutations). This was supported by significant congruence between topologies of the host phylogenetic tree and the symbiont community dendrogram (normalized Robinson– Foulds (nRF) test: Robinson-Foulds (RF = 0.97, *P* < 0.001)), although overall topological similarity between the trees was low. Together, these results support a weak but non-random phylosymbiotic pattern.

Only *Cladocopium* ITS2 type profiles exhibited a significant signal of cophylogeny with giant clam species (SS = 0.8976, R^2^ = 0.1024, *P* < 0.001; PACo, 999 permutations) (SFigure 11). No significant signal was detected for *Symbiodinium* (SS = 0.9805, R^2^ = 0.0195, *P* = 0.2982) or *Durusdinium* (SS = 0.9654, R^2^ = 0.0346, *P* = 0.1311). The strongest cophylogenetic relationships were observed between *T. maxima* and *Cladocopium* C1 lineages, and *T. derasa* and *Cladocopium* C93a lineages (SFigure 12a). *T. crocea* also exhibited low residual associations, although these were less consistent across lineages. *T. maxima* showed lower jackknifed squared residuals compared to other giant clam species, except *T. derasa*, indicating strongest contribution to the global phylogenetic signal (SFigure 12b).

## DISCUSSION

### Diversity of giant clam-associated Symbiodiniaceae

The Philippines is home to eight out of twelve giant clam species (bin Othman et al. 2010; Mies 2019). Most of these still occur naturally throughout the archipelago, except for the larger species like *T. gigas* and *T. derasa,* which are rare and considered locally extinct (Alcala 1986; Cabaitan and Conaco 2017; Dolorosa et al. 2024). Most *T. gigas* individuals found on Philippine reefs are likely the product of culture and restocking efforts that started in the 1980s (Mingoa-Licuanan and Gomez 2002; Gomez and Mingoa-Licuanan 2006; Moorhead 2018; Requilme et al. 2021). However, recent surveys in Palawan (Dolorosa et al. 2015; Mecha and Dolorosa 2020) and the Sulu Archipelago, southern Philippines (Muallil et al. 2024), have documented extant wild populations of *T. gigas*. The wide distribution of giant clam species across biogeographic regions of the Philippines provides a unique opportunity to investigate the dynamics of photosymbiont association with giant clams.

Philippine giant clams were associated with photosymbionts from the genera *Cladocopium, Durusdinium,* and *Symbiodinium*, mirroring symbiont communities of giant clam and coral hosts across the Indo-Pacific (Carlos et al. 1999; Deboer et al. 2012; Ikeda et al. 2017; Ravelo and Conaco 2018; Da-anoy et al. 2019; Lim et al. 2019; Mies et al. 2019; Morishima et al. 2019; Chao et al. 2021; Tañedo et al. 2021; Torres et al. 2021; Da-anoy et al, 2024; Guibert et al. 2025; Lee et al. 2025; Mies et al. 2025; Baquiran et al. 2026; Quijano et al. 2026). Most individuals in this study harbored all three photosymbiont genera, usually with one dominant genus and the others present only at background levels. Symbionts present at lower abundances may act as reserve populations for when environmental conditions favor a shift in photosymbiont composition (Mies et al. 2025).

*Cladocopium* was the most abundant genus associated with Philippine giant clams, with C93a, C1, C66, and C1c as the most prevalent ITS2 sequences. Of these, C66 and C1 are known to be broadly distributed in giant clams across the Indo-Pacific, with records from the South China Sea (Lim et al. 2019), Singapore (Lee et al. 2025), Vietnam (Mies et al. 2025), French Polynesia (Pochon et al. 2019), and the Great Barrier Reef (Weber 2009). C93a has been documented in the corals, *Porites* spp. (Grupstra et al. 2024) and *Acropora digitifera* (Buzzoni et al. 2026), in Palau, as well as in soritid foraminiferans in Guam (Pochon et al. 2007). C1c has so far only been detected in corals (Quigley et al. 2020; Liang et al. 2021).

The prominent *Durusdinium* sequences in giant clams were D1, D4, D5, and D9b. These sequences constitute the D1-D4 profile defining *D. trenchii* (LaJeunesse et al. 2014) and the D4/D5/D9b radiation. *D. trenchii* and other D1-radiation photosymbionts have been detected in *T. noae* from the South China Sea and the West Indo-Pacific (DeBoer et al. 2012; Lim et al. 2019), as well as in the Red Sea (Rossbach et al. 2021). The D4/D5/D9b radiation has been reported in *T. noae* and *T. squamosa* in Singapore (Lee et al. 2025).

*Symbiodinium* sequences A3 and A6b were present in all Philippine giant clam species but at low abundance. A3 is affiliated with *Symbiodinium tridacnidorum*, which is prevalent in Indo-Pacific giant clams. In our study, A3 was significantly associated with *H. hippopus*, which is also the species from which this symbiont was first isolated (Lee et al. 2015). A6b, on the other hand, appears to be a relatively minor ITS2 sequence with limited records in corals such as *Platygyra daedalea* and *Acropora digitifera* from the South China Sea (Jiang et al. 2021; Samshuri et al. 2023).

### Drivers of giant clam symbiont community structure

#### Host specificity and flexibility

Photosymbiont communities associated with giant clams are structured by the host. The dominant symbiont in each species was generally similar to observations in giant clams from other regions. For example, *Cladocopium* is also dominant in *T. crocea*, *T. gigas*, *T. maxima,* and *H. porcellanus* from various sites within the Indo-West Pacific (Baillie et al. 2000; LaJeunesse 2001; Weber 2009; DeBoer et al. 2012; Lim et al. 2019; Pochon et al. 2019), while *Durusdinium* is prevalent in *T. noae* from the Dongsha Atoll in the northern South China Sea (Lim et al. 2019). There are, however, some exceptions. In particular, Philippine *T. squamosa* samples were dominated by *Durusdinium*, which is in contrast to other Indo-Pacific *T. squamosa* populations that host mostly *Cladocopium* or *Symbiodinium* (Weber et al. 2009). In addition, Philippine *T. maxima* hosted *Cladocopium*, whereas individuals from the Red Sea (Rossbach et al. 2021) and various sites in the Indo-Pacific region (Weber 2009) were found to be *Symbiodinium*-dominated. These variations in symbiont community may reflect differences in local reef environments and in the available photosymbiont pool.

Giant clams acquire symbionts horizontally from the environment, the surrounding seawater, sediments, or even fecal pellets from neighboring giant clams (Trench et al. 1981; Morishima et al. 2019). Thus, the local symbiont pool will influence initial colonization and may be reflected in the composition of the community in the giant clam (Fitt and Trench 1981; Hirose et al. 2006). This is most apparent during early ontogeny, when veliger larvae first start to acquire photosymbionts (Fitt et al. 1984; Mies et al. 2018; Long et al. 2021). As veligers develop, winnowing of the photosymbiont community occurs (Ikeda et al. 2017). Although the mechanisms by which this happens remains unknown, they possibly involve selectivity of host receptors (Mohamed et al. 2016; Yoshioka et al. 2023), selection by the microenvironment in the host (Holt et al. 2014; Mies et al. 2025), or competitive interactions within the symbiont community (Abbott et al. 2021). Ultimately, the most well-adapted symbionts will proliferate and eventually become the dominant members of the community, while other symbionts remain in the background.

The degree of specificity or flexibility of symbiont associations in giant clams could be driven by differences in host traits and metabolic requirements, as these influence the microenvironments experienced by the photosymbionts. Giant clams likely associate with symbionts that can conduct photosynthetic activity and provide optimal nutrient exchange under the habitat conditions preferred by the host. For example, *T. noae* typically occurs in shallower reef areas where temperature and light levels may fluctuate widely, whereas *T. gigas* is found in more sheltered, sunlit reefs, and *T. squamosa* can thrive in deeper habitats (Rosewater, 1965; Jantzen et al. 2008; Watson and Neo, 2021). Burrowing species like *T. crocea* and *T. maxima* may be partially protected from external factors by hard substrates (Toonen et al. 2012; Chung et al. 2026), whereas *H. hippopus* and *H. porcellanus,* which are typically found on loose, sandy substrates may be impacted by wave activity (Panggabean et al. 2006; de Guzman et al. 2023). While host-specific mantle coloration and patterning may protect photosymbionts from UV-radiation or help improve photosynthetic efficiency through wavelength conversion (Holt et al. 2014; Rossbach et al. 2020), differences in temperature, light, and even wave action in the giant clam habitat will influence the photosynthetic output of symbionts and challenge their tolerance for environmental fluctuations. Importantly, giant clams differ in their dependence on nutritional contributions from photosymbionts (Klumpp et al. 1992; Klumpp and Griffiths 1994). A comparison across species revealed that *T. gigas* and *T. derasa* are relatively more dependent on autotrophic sources of nutrition compared to other giant clams (Guibert et al. 2025). These species may therefore prefer symbionts with better photosynthate translocation capacity, such as *Cladocopium*, which is known to be more efficient at sharing nutrients with the host compared to *Durusdinium* (Cantin et al. 2009; Allen-Waller and Barott 2023), although there are exceptions (Kemp et al. 2023; Turnham et al. 2023). Nevertheless, each giant clam species exhibits some flexibility in their symbiotic associations, partnering with different symbiont genera, or with different strains of the same genus. This could reflect a strategy that balances metabolic complementation with environmental tolerance. Functional differences in symbiotic interactions could contribute, in part, to the observed distribution of giant clam species, where *T. maxima* and *T. squamosa* are most widespread, followed by *T. gigas*, *T. derasa*, *T. noae*, *T. crocea* and *H. hippopus* with intermediate range, and *H. porcellanus* found only in a few areas (bin Othman et al. 2010; Neo et al. 2017; Mies et al. 2019).

#### Environmental drivers

Environmental factors further contribute to structuring of giant clam-associated symbiont communities. In archipelagic systems like the Philippines, water currents and basin structure limits connectivity among reefs (Pata and Yñiguez 2019; 2021). If dispersal is limited for free-living photosymbionts (Howells et al. 2013; Baums et al. 2014; Wepfer et al. 2020), distinct photosymbiont pools are largely retained within specific areas, explaining the general variation in Symbiodiniaceae communities across the 11 different reef sites and five biogeographic regions sampled in this study. This provides a plausible cause for the region-specific photosymbiont types observed within some host species, while still allowing general overlap among hosts exposed to similar environments. For example, the biogeography-driven structuring of Symbiodiniaceae that we observed in this study mirrors spatial genetic structuring of *T. crocea* populations from the North Philippine Sea and South Philippine Sea (Ravago-Gotanco et al. 2007), suggesting the presence of a dispersal boundary. Additionally, the composition of other co-occurring giant clam species and photosymbiotic organisms (e.g., corals, sponges, foraminiferans) in different reef areas may influence the availability of Symbiodiniaceae through release and exchange of photosymbionts within reef environments. Further studies to characterize photosymbionts across diverse reef areas, as well as syntopic taxa hosting Symbiodiniaceae, may help disentangle region-specific differences in photosymbiont communities associated with giant clams.

Giant clam symbiont communities are also influenced by factors such as temperature (Leggat et al. 2003; Armstrong et al. 2022), light availability (Adams et al. 2013; Liu et al. 2020), and water quality (Toonen et al. 2012; Lee et al. 2024). Of these, temperature is one of the stronger determinants of host-associated Symbiodiniaceae composition. Although not as strong as host-driven structuring, Symbiodiniaceae communities also varied by broad thermal regimes. However, instead of thermotolerant *Durusdinium* symbionts, we found that *Symbiodinium* A6b and A3g are prevalent in the warmest thermal regime while co-occurring with A3 (*S. tridacnidorum*) and *Cladocopium* C66. *Symbiodinium tridacnidorum* is considered a heat-sensitive species (Cumbo et al. 2018). This same species was dominant in *T. maxima* and *T. squamosa* from the Red Sea, a region that also experiences elevated temperatures (Pappas et al. 2017), suggesting local adaptation. Unexpectedly, we detected *Durusdinium* ITS2 sequences in the thermal regime with the lowest temperature range, although this signal is likely attributed to overrepresentation of *T. noae* individuals. The close association of this particular giant clam species with *Durusdinium* may contribute to its success in the shallower reef areas (Neo et al. 2018) where dynamic fluctuations in temperature are common. Taken together, these findings indicate that temperature alone may be insufficient to explain the observed patterns in giant clam-symbiont associations, as these trends may be influenced by a combination of multiple abiotic and biotic factors, highlighting the need for deeper investigations.

#### Symbiont community dynamics

Two decades after translocation from the Silaqui ocean nursery in Bolinao to various reef sites in the Philippines, *T. gigas* individuals still retained a similar symbiont community dominated by closely-related photosymbionts from the genus *Cladocopium*. This suggests that symbionts that were initially established during development are maintained even after prolonged residence in a new environment, which may partly explain the low symbiont diversity observed in *T. gigas*. Shifts in dominance of closely-related symbionts, especially in individuals belonging to the same haplotype, may have resulted from stochastic events or were triggered by environmental stressors that resulted in some symbiont loss, followed by proliferation of pre-existing background symbionts or colonization by symbionts from the environment. Similar environmental structuring of ITS2 types has been documented in *T. maxima* across the Red Sea, where photosymbiont profiles varied along an environmental gradient (Rossbach et al. 2021), and may reflect processes comparable to photosymbiont shuffling reported in heat stressed *T. squamosa* (Mies et al. 2025).

Notably, *T. gigas* individuals from Palawan, which mostly belonged to different COI haplotypes, also hosted symbionts that differed from the majority of restocked *T. gigas*. This suggests that the Palawan individuals could have originated from culture cohorts that used different parental broodstocks or are natural recruits that acquired and maintained a different set of symbionts from the local environment. Juvenile natural recruits in Camiguin also exhibited different symbionts from translocated clams, with C1 as the dominant DIV, supporting the idea that early symbiosis in giant clams is influenced by local photosymbiont availability. This observation also suggests that *T. gigas* prefers to associate with members of the *Cladocopium* lineage. The influence of host ontogeny or duration of exposure to local environmental conditions on the establishment and stability of giant clam-Symbiodiniaceae partnerships is expected due to the horizontal acquisition of photosymbionts (Fitt et al. 1984). Nevertheless, because younger individuals and natural recruits were underrepresented in this study, further transplantation experiments that explicitly include early life stages and track photosymbiont profiles through time are needed to disentangle the relative roles of host age and environmental filtering in giant clam-Symbiodiniaceae associations. Furthermore, it would be of interest to examine how the symbiont communities in other giant clam species with variable symbiont specificities would behave when translocated to different environments.

### Phylosymbiosis and cophylogeny in giant clams and their symbionts

Phylosymbiosis and cophylogeny link host evolutionary relationships with symbiont community diversity. The presence of phylogenetically structured associations between giant clams and their photosymbionts is notable, given that giant clams exclusively acquire their photosymbionts through horizontal transmission. However, it is plausible that giant clams, while initially taking up a broad range of Symbiodiniaceae, subsequently exert selective pressures through physiological compatibility, host-mediated regulation (Armstrong et al. 2018; Pang et al. 2022), and host recognition (Li et al. 2025; Yi et al. 2025). Moreover, symbionts may also exhibit preferences for certain host genetic backgrounds or traits (Johnston et al. 2022; Li et al. 2025). Over evolutionary timescales, repeated post-acquisition filtering mechanisms may promote consistent associations between particular host taxa and photosymbiont lineages, giving rise to the observed patterns of phylosymbiosis and cophylogeny despite horizontal transmission (Lim and Bordenstein 2020). It is likely that preferred host-symbiont partnerships impart functional benefits to the giant clam holobiont, such as efficient nutrient transfer or adaptability to habitat conditions, although further investigations are needed to test these ideas.

### Conclusions

Our study revealed that Philippine giant clams harbor diverse Symbiodiniaceae communities. The composition of these communities is shaped by complex interactions between host species or genotype and environmental factors, such as temperature, light, water quality, and available symbiont reservoirs, that vary across different spatial scales, as well as symbiont community dynamics through processes such as colonization, competition, and succession. Some giant clam hosts, such as *H. hippopus*, *T. maxima*, and *T. crocea*, show greater flexibility in the types of symbionts that they associate with, suggesting adaptability to a wider range of environmental conditions. On the other hand, *T. gigas* appears to associate mostly with members of *Cladocopium*. The choice of symbiotic partners may impact metabolic exchange within the holobiont and, in part, influence adaptability and distribution of species. Giant clams with broader symbiont preference could thrive in a wider range of environments, whereas those with narrower preference may be more restricted in distribution and could potentially be more vulnerable to environmental perturbations that result in loss of symbionts. To further clarify the mechanisms governing giant clam-Symbiodiniaceae associations, particularly how host identity overrides influence of environment in an organism that horizontally transmits its photosymbiont, further studies incorporating broader geographic sampling, genomic comparisons, and controlled translocation and symbiont inoculation experiments are warranted. Ultimately, as coral reefs face escalating thermal stress, characterizing the factors that govern photosymbiont diversity and flexibility in giant clams will be crucial to understanding their resilience and adaptive potential under climate change.

## Data availability

Raw sequence reads can be accessed from the NCBI Short Read Archive database under BioProject PRJNA749183. The datasets we generated and analyzed are available on Figshare (https://doi.org/10.6084/m9.figshare.33004178) – raw sequences, processed data, COI sequences. The scripts used for data processing and analysis are available on GitHub (https://github.com/jbquijano/001_gcla).

## Acknowledgements

We thank the local government units of Alaminos, Bolinao, and Anda, Pangasinan; Mabini and Bauan, Batangas; Masinloc, Zambales; Vinzons, Calaguas, Camarines Norte; Sablayan, Occidental Mindoro; Panglao and Calape, Bohol; Guiuan, Samar; Puerto Princesa, Palawan; and Guinsiliban, Camiguin for supporting the field surveys. We thank Dr. Ronnie Estrellada for generously contributing materials from the Semirara Marine Hatchery Laboratory. We thank Lala Grace Calle, Ian Joseph De Guzman, Eymard John Sy, Keana Tan, Michael Angelou L. Nada, Ronald de Guzman, Ben Jack Gabuay, and other staff of Bolinao Marine Laboratory for their invaluable assistance in the field surveys and sample collection. This study was funded by grants from the Philippine Council for Agriculture, Aquatic, and Natural Resources Research and Development of the Department of Science and Technology to PCC, RGD, and CC (QMSR-MRRD-MEC-295-1449; QMSR-MRRD-MEC-314-1542/1543/1544/1545).

## Contributions

JBQ: data curation; formal analysis; methodology; visualization; writing – original draft preparation; writing – review & editing

KMT: investigation; methodology; writing – review & editing

JIPB: formal analysis; writing – original draft preparation; writing – review & editing

GJM: investigation; writing – review & editing

JNR: investigation; writing – review & editing SLS: investigation; writing – review & editing

RGD: investigation; writing – review & editing

PCC: conceptualization; funding acquisition; investigation; writing – review & editing

CC: conceptualization; funding acquisition; investigation; formal analysis; supervision; writing – original draft preparation; writing – review & editing

