## Supplementary figures for "Host and environment shape the giant clam-associated photosymbiont community"

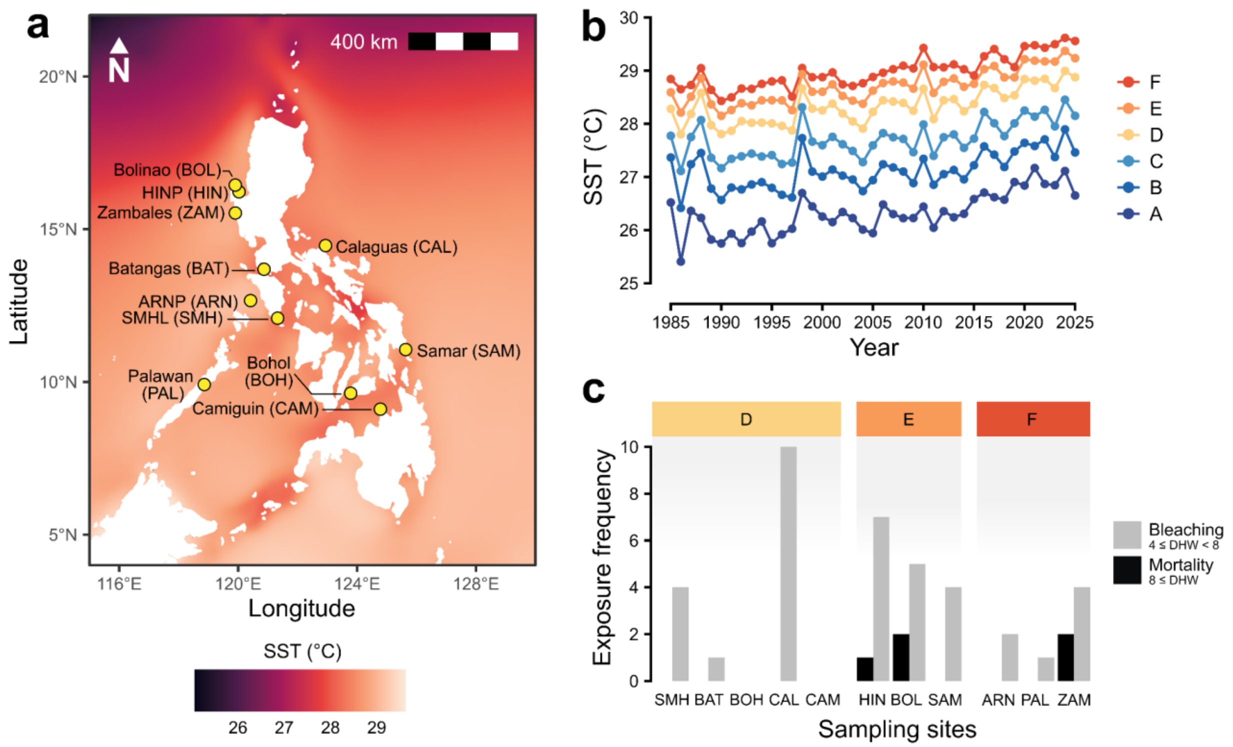


**SFigure 1. Thermal profile of the Philippine seas.** (**a**) Mean sea surface temperatures (SST) across the Philippines from 1985 to 2025. (**b**) Annual mean SSTs among the identified thermal regimes. (**c**) Frequency of exposure presented as number of years with bleaching- (gray) and mortality-inducing (black) degree heating weeks (DHWs).

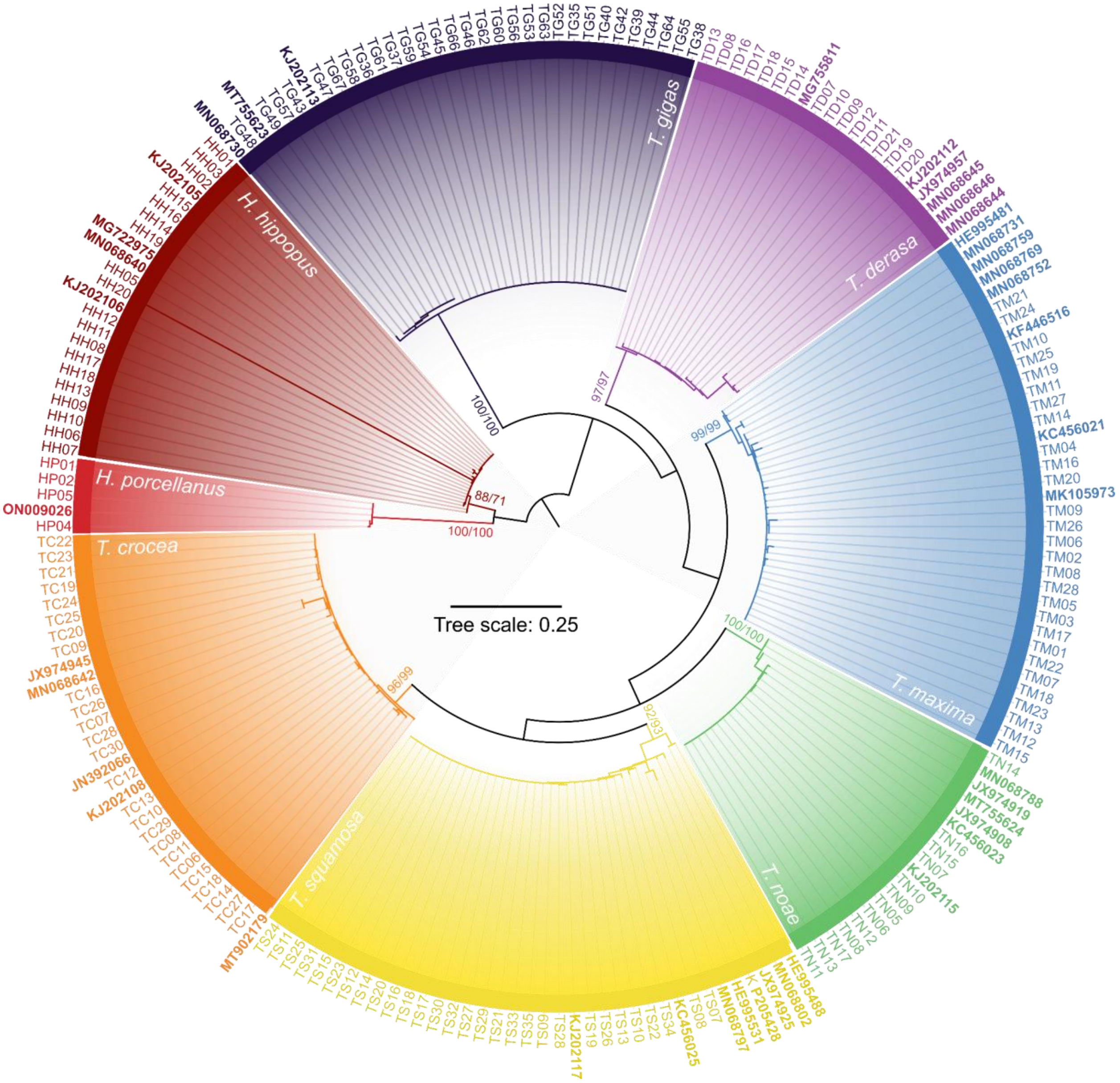


**SFigure 2.** Phylogeny of sampled giant clam individuals based on the COI gene. Branch support values are shown as SH-aLRT and UFBoot percentages at each clade. Branches are colored by species. Tips in bold indicate reference sequences from NCBI. Scale bar represents branch length.

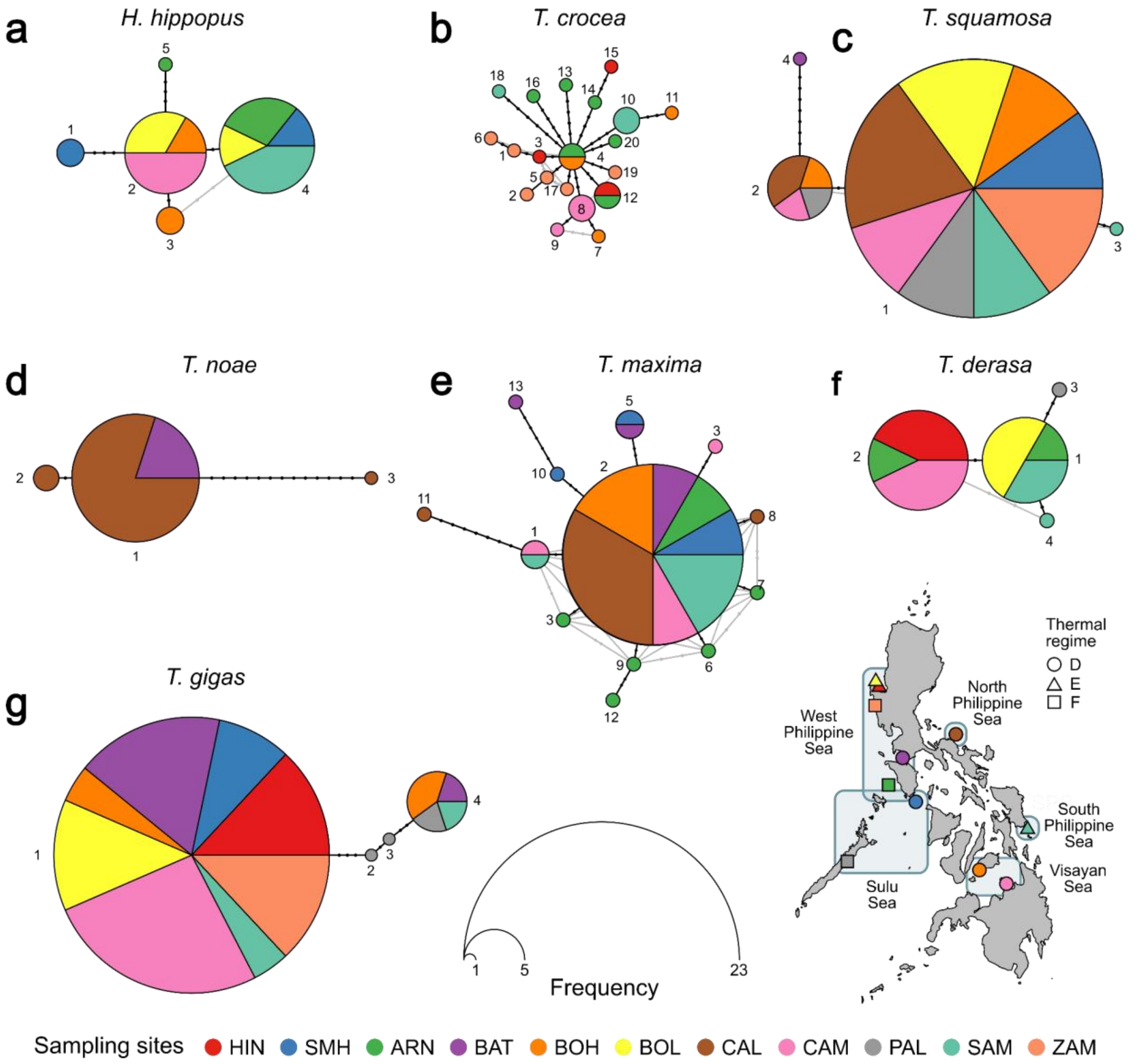


**SFigure 3. COI haplotype networks for giant clam species.** Haplotypes detected in *H. hippopus* (**a**), *T. crocea* (**b**), *T. squamosa* (**c**), *T. noae* (**d**), *T. maxima* (**e**), *T. derasa* (**f**), and *T. gigas* (**g**). Only one haplotype was identified for *H. porcellanus* (not shown). Each numbered node on the network represents a unique haplotype for each species. Node size is proportional to haplotype frequency. Dots along connecting lines indicate mutational steps between haplotypes. Pie colors represent sampling sites. The map indicates sites, biogeographic regions (highlighted areas), and thermal regimes (marker shapes).

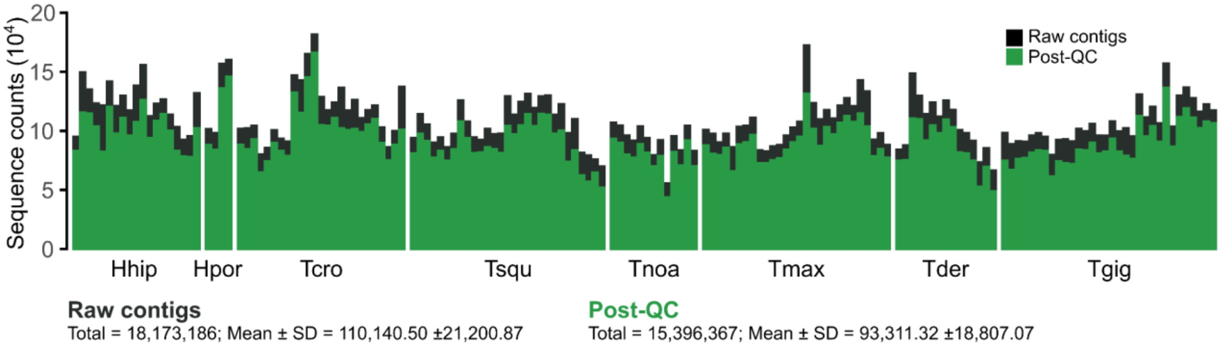


**SFigure 4. Symportal run statistics.** Sequence counts before (black) and after (green) quality control and filtering.

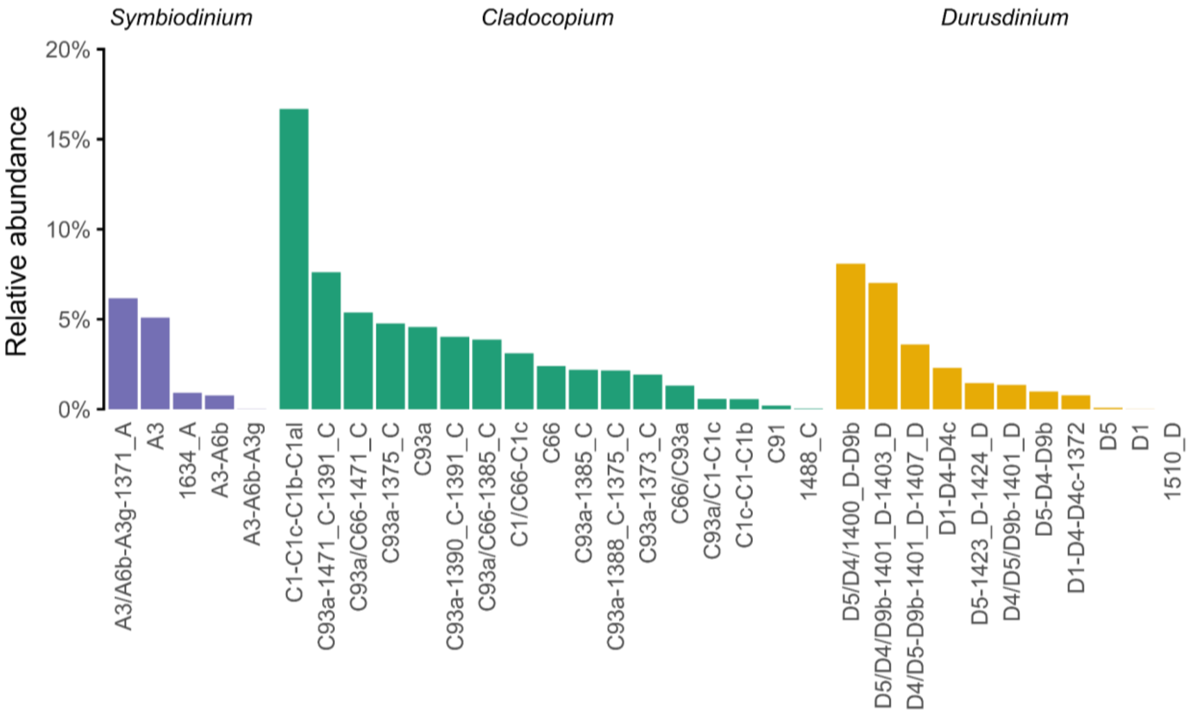


**SFigure 5. Relative abundance of ITS2 type profiles associated with Philippine giant clams**.
Bars are grouped by symbiont genus.

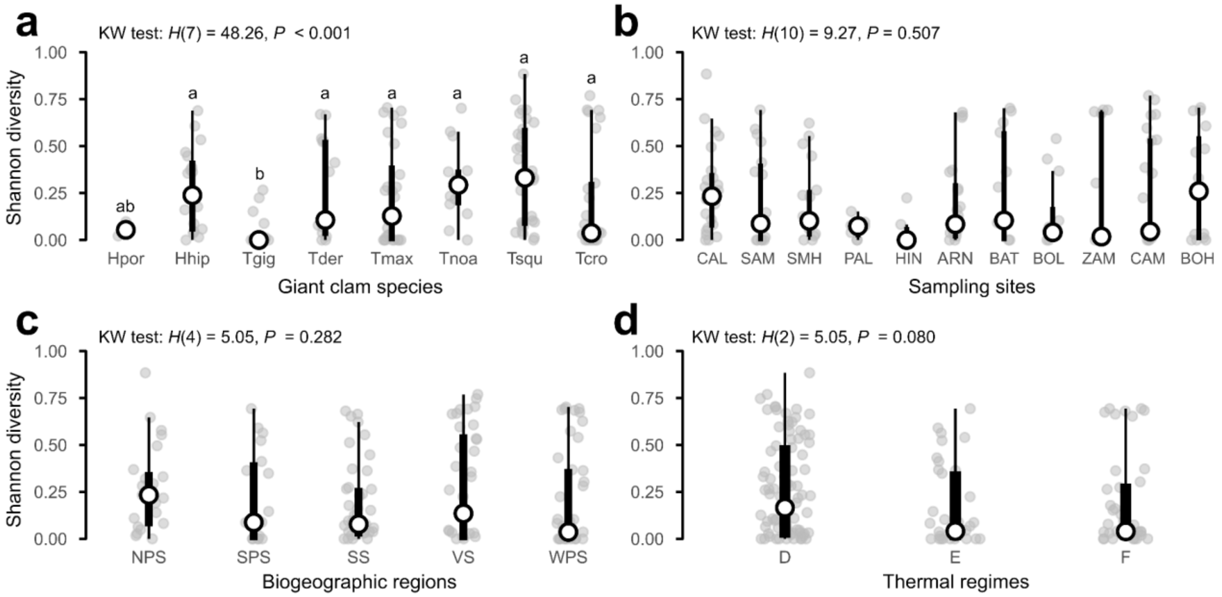


**SFigure 6. Symbiodiniaceae diversity across groups.** Box plots of Shannon diversity of Symbiodiniaceae communities among giant clam species (**a**), across sampling sites (**b**), biogeographic regions (**c**), and thermal regimes (**d**). Large white circles indicate medians, and background points represent individual samples. Kruskal-Wallis test results are shown above each plot.

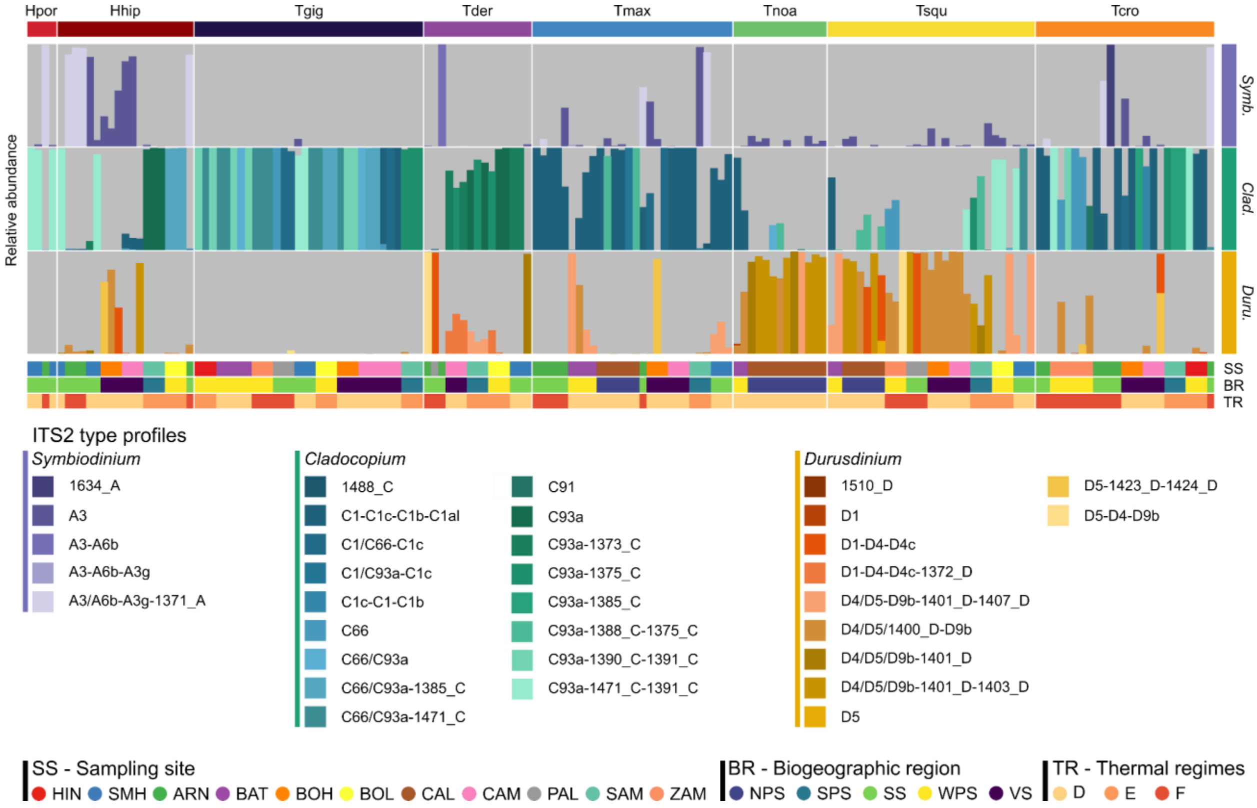


**SFigure 7. ITS2 type profile composition across groups.** Relative abundance of ITS2 type profiles in each giant clam individual. Colors denote ITS2 type profiles. The top color bar indicates host species. The right color bar indicates symbiont genus. The bottom color bars indicate sampling site, biogeographic region, and thermal regime.

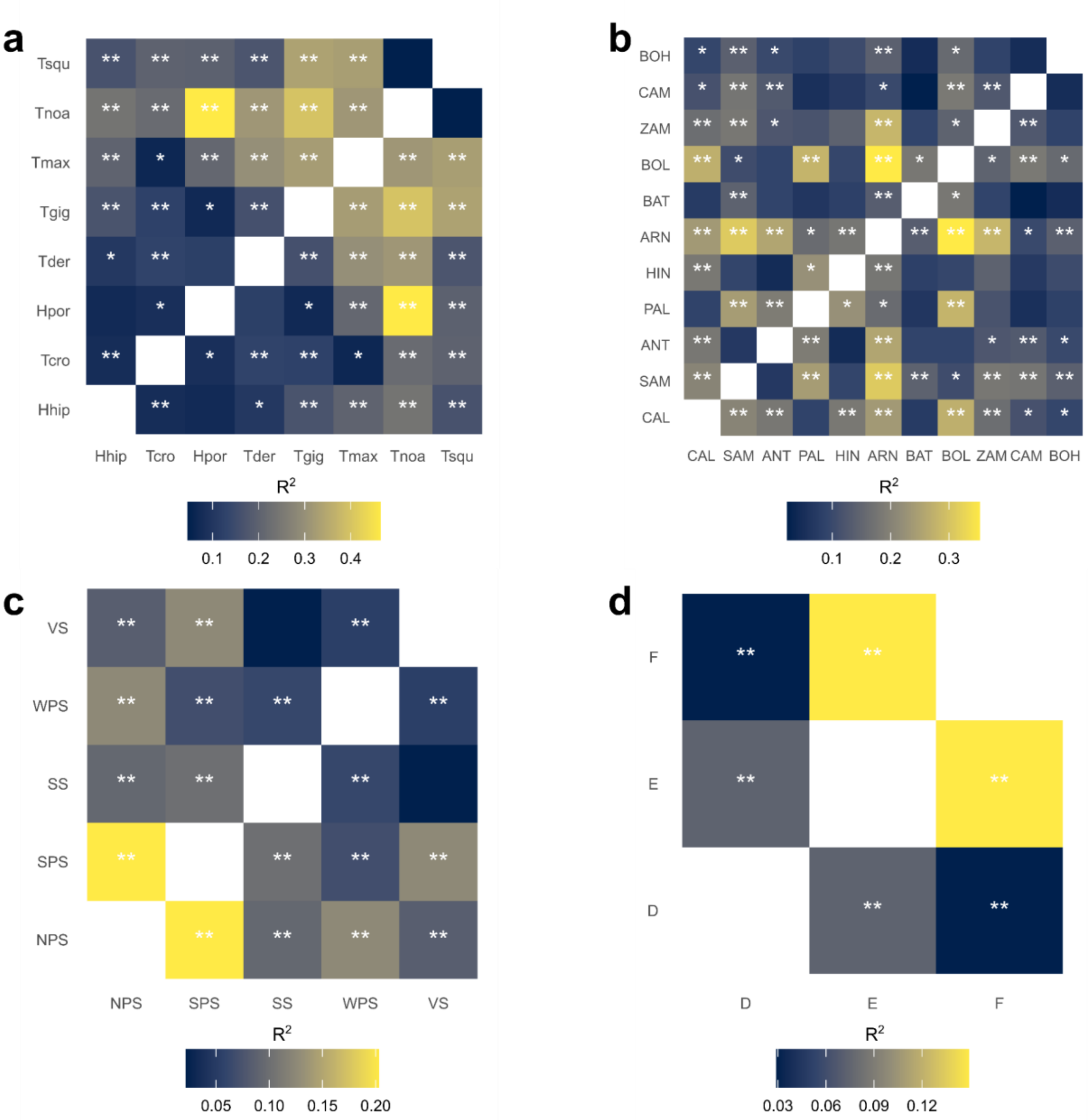


**SFigure 8. Pairwise PERMANOVA comparisons of Symbiodiniaceae community composition.** Pairwise comparisons of Symbiodiniaceae communities based on ITS2 sequences across species (**a**), sampling sites (**b**), biogeographic regions (**c**), and thermal regimes (**d**). Significant tests are indicated by asterisks ($Q \leq 0.05$ *, $Q < 0.01$ **, $Q < 0.001$ ***). The color gradient denotes pairwise $R^2$ values.

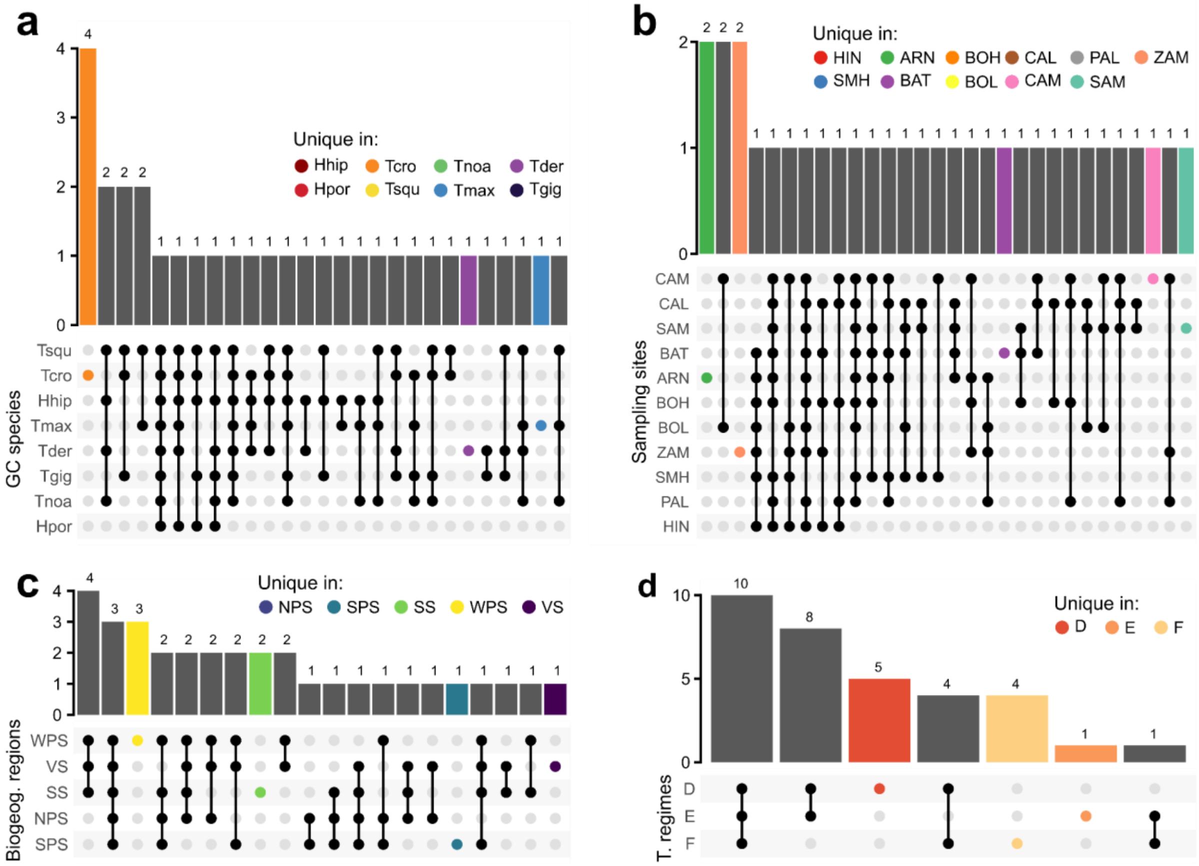


**SFigure 9. Shared and unique ITS2 type profiles across groups.** UpSet plots showing shared and unique ITS2 type profiles among species (**a**), sampling sites (**b**), biogeographic regions (**c**), and thermal regimes (**d**). Bars indicate the number of ITS2 type profiles per group combination (connected dots). Colors denote ITS2 type profiles unique to a group.

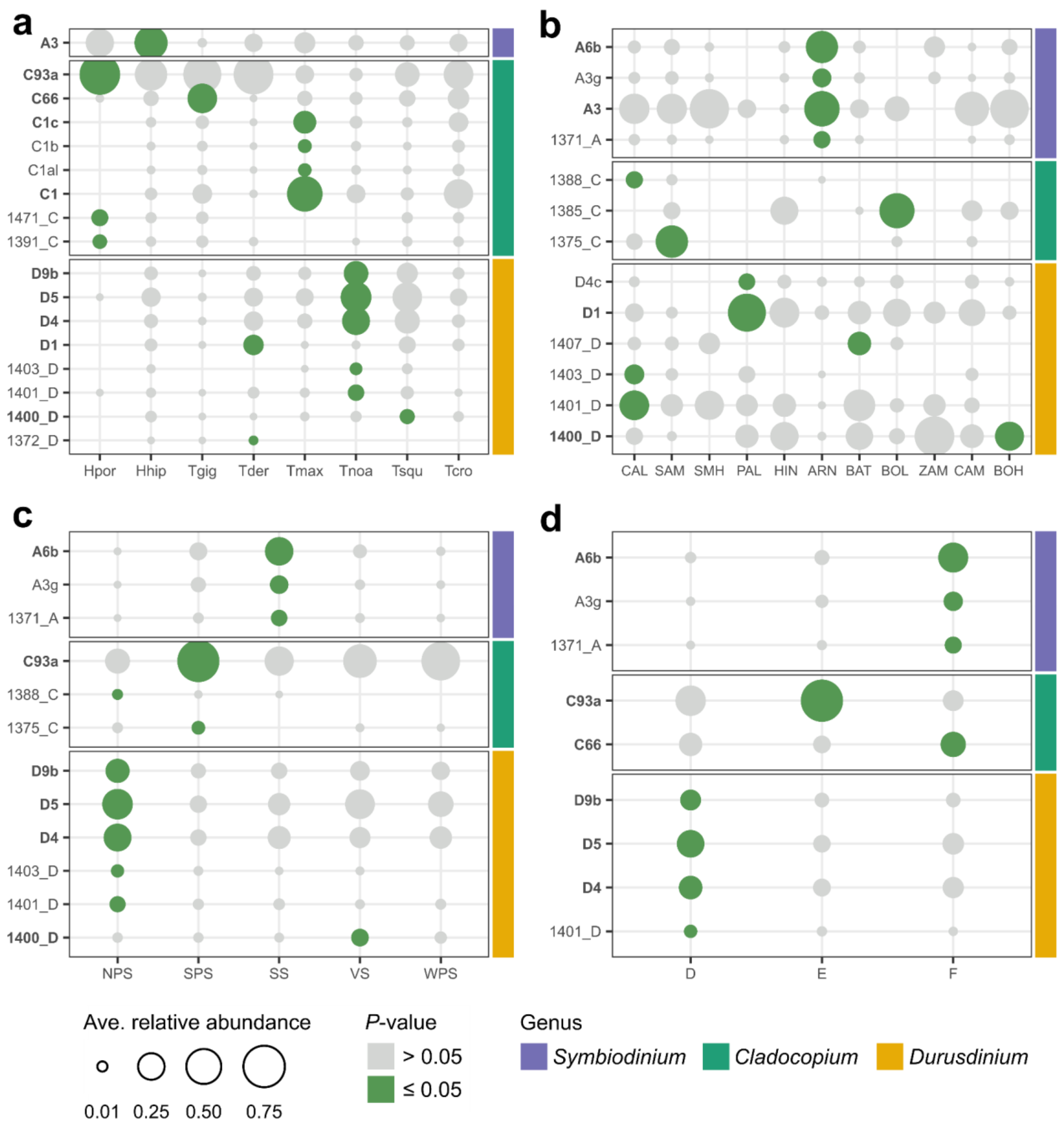


**SFigure 10. Defining intragenomic variants (DIVs) associated with specific groups.** Indicator species analysis was used to identify ITS2 DIVs significantly associated ($P \leq 0.05$, IndVal $\geq 0.5$) with species (**a**), sampling sites (**b**), biogeographic regions (**c**), and thermal regimes (**d**). Dot size represents the average relative abundance of each DIV per group. DIVs are separated and colored by Symbiodiniaceae genus. Dot colors represent *P*-value (green, *P*-value < 0.05; gray, *P*-value ≥ 0.05). Major DIVs are highlighted in bold.

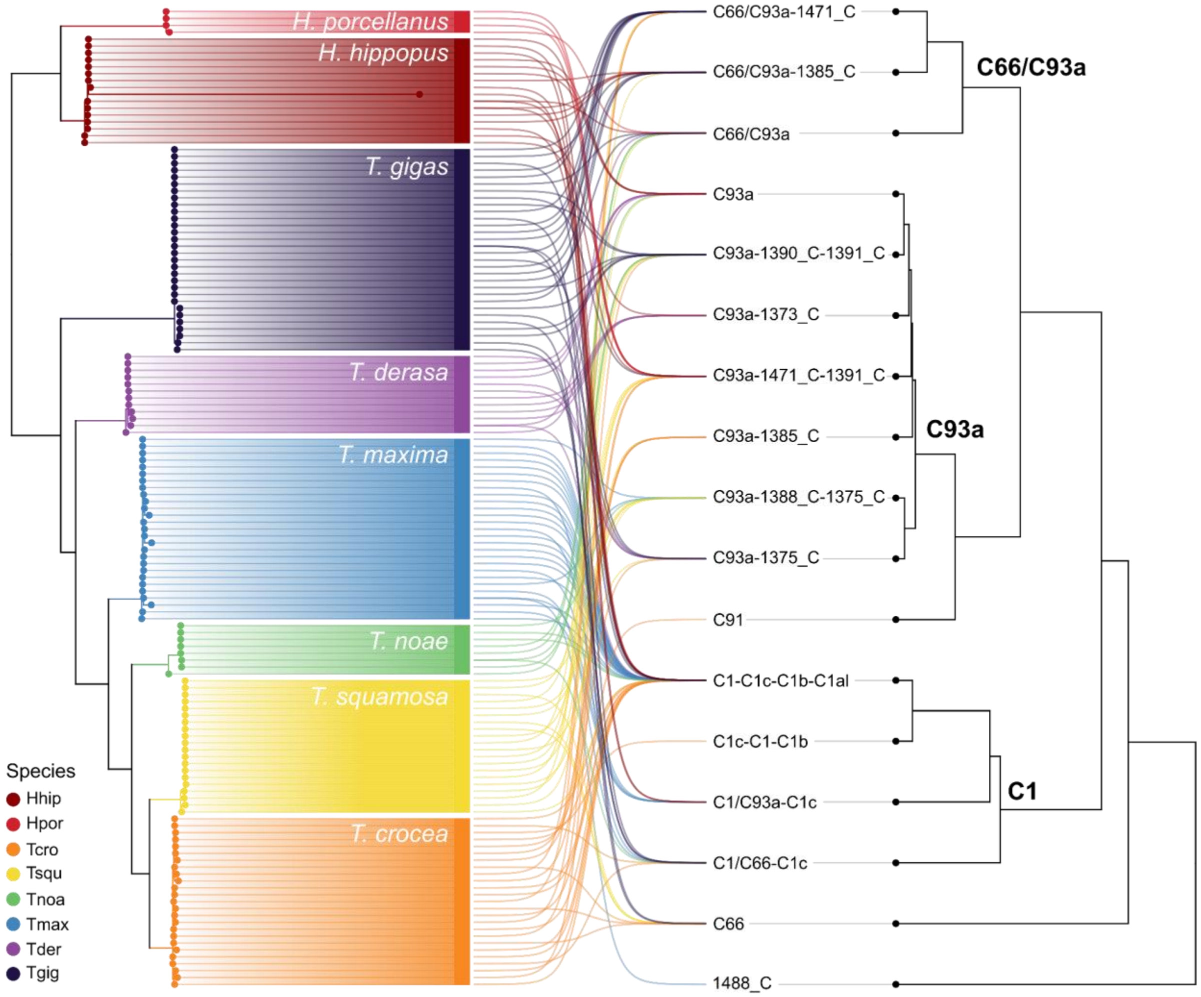


**SFigure 11. Cophylogeny of giant clam species and *Cladocopium*.** Tanglegram depicts the associations between 142 giant clam individuals and 17 *Cladocopium* ITS2 type profiles based on PACo analysis. Host phylogeny was reconstructed from COI sequences using IQ-TREE, while the symbiont dendrogram was generated using UPGMA based on UniFrac distances derived from defining intragenomic sequence variants (DIVs). Leaves and association links are colored by giant clam species. Symbiodiniaceae clades are labeled by the most dominant DIV.

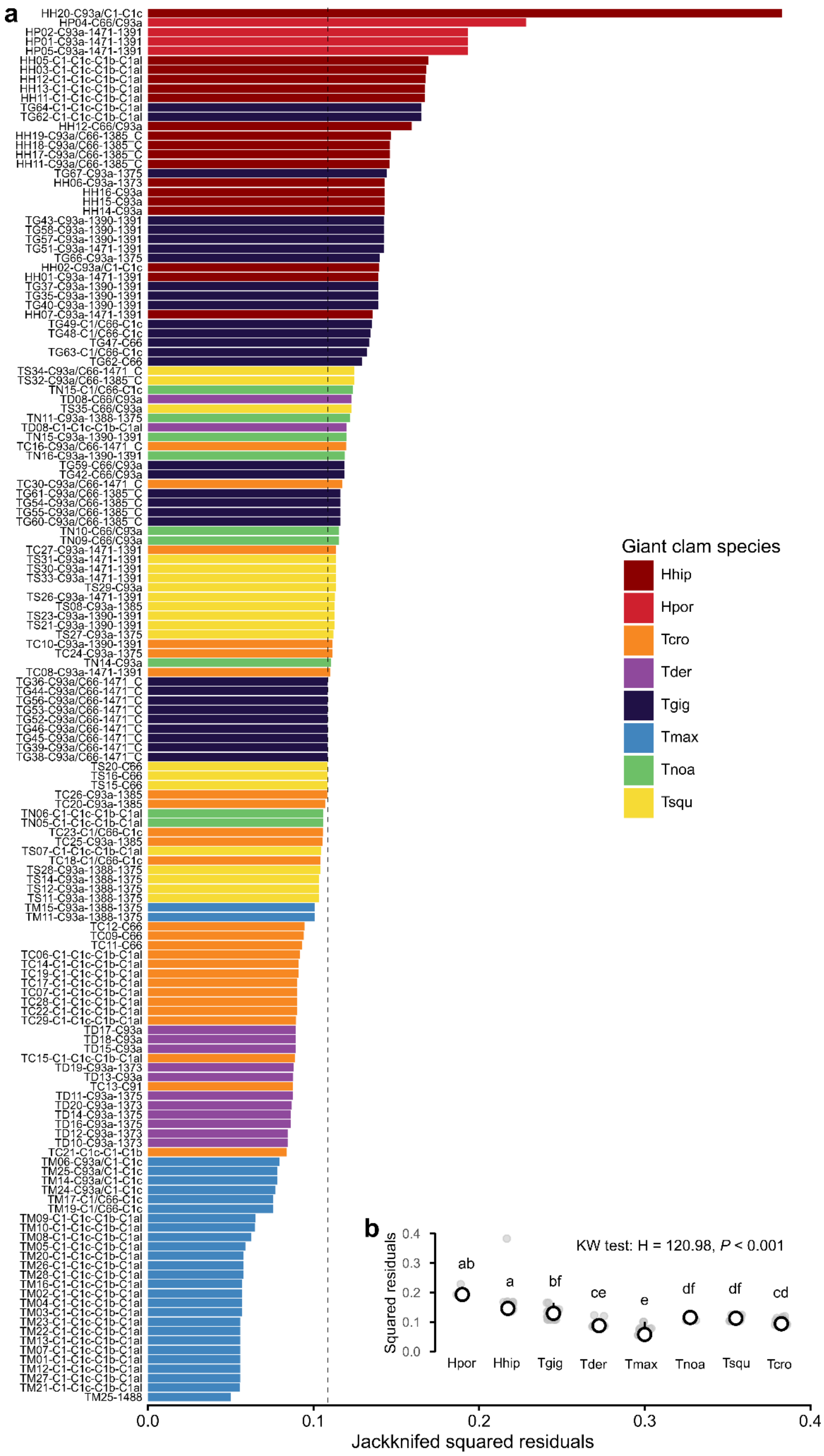


**SFigure 12. Host-symbiont link contributions to cophylogenetic fit inferred from PACo.** (**a**) Jackknifed squared residuals derived from PACo representing the contribution of each giant clam-

ITS2 type profile link to the global cophylogeny fit. The dashed line represents median squared residual. (**b**) Comparison of jackknifed squared residuals among host species. Different letters indicate significant differences among species based on Dunn's post-hoc test following a Kruskal-Wallis test ($P < 0.05$).
